# Modeling the metabolic heterogeneity of high-grade serous ovarian cancer solid tumors in 3D Microphysiological systems

**DOI:** 10.64898/2026.06.30.735360

**Authors:** Paula M. Mañán Mejías, Nicha Boonpattrawong, Michael Bérubé, Ellinore K. Letts, Finnbar Reed-McBain, Ana Sofía Peraza Muñuzuri, Yalmarie Numan Vázquez, Manish Patankar, María Virumbrales-Muñoz

## Abstract

High-grade serous carcinoma (HGSOC) is the deadliest subtype of ovarian cancer, characterized by high metastatic rates. HGSOC is typically diagnosed at late stages, and treatment options are limited, resulting in a 60% recurrence rate. HGSOC cells exhibit metabolic plasticity, dynamically shifting between glycolysis and oxidative phosphorylation (OXPHOS) to meet energy demands for tumor progression. To evaluate therapeutic strategies that target metabolic vulnerabilities, we developed a microphysiological system (MPS) that recapitulates the heterogenous cell states and bioenergetic distribution of HGSOC solid tumors. Our platform utilized HGSOC spheroids embedded in a collagen hydrogel that mimics the extracellular matrix to capture tumor progression in the ovary. We used atovaquone (ATO), an FDA-approved OXPHOS inhibitor, to prototype the capabilities of our platform to investigate metabolic plasticity in HGSOC. Treatment with ATO decreased viability and invasion of HGSOC spheroids. Crucially, ATO exhibited no cytotoxicity toward biomimetic blood vessels, preserving their integrity and permeability. Metabolic imaging revealed that ATO induces an oxidative state in the outer region of the spheroids. At the invasive front, ATO disrupted mitochondrial organization, forcing collective cell migration and eventually inducing breakdown of mitochondrial networks. Furthermore, ATO decreased YAP/TAZ pathway activity in the outer region of the spheroid, providing a potential mechanism for hindered cell invasion. Collectively, our data demonstrates that a low-potency OXPHOS inhibitor like ATO can effectively target metabolic plasticity to suppress HGSOC spheroid progression. Overall, this platform successfully recapitulated metabolic heterogeneity and provided a workflow for safely testing other drugs that target cancer metabolism.

## Introduction

High-grade serous ovarian carcinoma (HGSOC) is the most lethal gynecological malignancy^1^. Debulking surgery plus platinum-based chemotherapy remains the standard of care, yet there is frequent recurrence after these treatments^2^, and eligibility for targeted therapies (e.g., PARP inhibitors) is limited. The effectiveness of these therapy regimens long-term is limited, with only 30-40% of patients with advanced stage HGSOC surviving for five years or more after initial diagnosis^3^. Therefore, there is an urgent need to develop novel therapies for effective treatment of HGSOC, that improve outcomes for these patients.

Dysregulated cellular bioenergetic pathways in tumors provide significant opportunities for developments of novel cancer therapeutics ^4^. While it is generally accepted that cancer cells switch to an aerobic glycolysis phenotype^5^, many studies have now confirmed that OXPHOS is active in cancer cells (including in ovarian cancer)^6^, and that tumors are metabolically heterogeneous^7–9^. Practically, this means that cancer cells are capable of metabolic plasticity, where they can switch their utilization of metabolic pathways to adaptively fulfill bioenergetic demands for processes required for tumor progression: cell proliferation, migration and survival^7, 10^. Furthermore, ovarian cancer cells can switch from an OXPHOS-low to an OXPHOS-prevalent phenotype as they acquire chemoresistance to platinum agents, which has been reversed *in vitro* using OXPHOS inhibitors^11^, underscoring a therapeutic opportunity for targeting metabolic plasticity^12^. Notably, all these processes are fundamentally affected by a 3D microenvironment^13^. For example, 3D cancer cell migration requires more energy than in 2D^14^, since cells must constantly adapt their cytoskeleton and secrete enzymes to remodel their surrounding ECM to break through its physical constraints^15, 16^. As such, mitochondrial network remodeling and subcellular location in the cell cytoplasm becomes highly important, to provide cells with the necessary energy at their leading edge. However, the prevalence of OXPHOS-high vs OXPHOS-low metabolisms during cell migration processes remains incompletely understood.

A signaling pathway that has recently emerged as a key regulator of metabolic plasticity is the Hippo pathway^17^. This pathway acts as a critical sensor, responding to diverse microenvironmental cues (e.g., mechanical crowding, inflammation, hypoxia, metabolic stress) by activating adaptive transcriptional programs. Specifically, high expression of the transcription factor Yes-associated protein 1 (YAP1) and co-regulator TAZ (Transcriptional coactivator with PDZ-binding motif) in HGSOC is linked to the upregulation of cell survival genes and chemoresistance^18^. When cells in healthy tissues find themselves in dense and nutrient-depleted 3D environments, YAP/TAZ activity is normally suppressed (via AMPK phosphorylation leading to degradation) to prevent cell growth when conditions are unfavorable^19, 20^. However, cancer cells can aberrantly activate the YAP/TAZ pathway, allowing YAP1 and TAZ to translocate into the nucleus and promote cell proliferation, stemness and survival^21^. Thus, we posit that therapeutic OXPHOS inhibition may induce metabolic stress and lead to YAP1 degradation and therefore halt progression.

Given the therapeutic potential of reprogramming the YAP/TAZ pathway, considerable efforts have focused on targeting the OXPHOS-YAP1 as a therapeutic strategy across multiple cancer types^20^. However, the ubiquitous nature of OXPHOS and its high metabolic efficiency imposes limitations on this concept. Systemic inhibition frequently results in narrow therapeutic windows and dose-limiting toxicities.^22, 23^. Atovaquone (ATO), an FDA-approved antimalarial therapy and OXPHOS inhibitor, has recently risen to the spotlight as a potential alternative therapy in HGSOC and other cancers. Specifically, ATO has demonstrated to specifically target cancer cells and delay cancer growth in established cell lines and mouse models. Interestingly, the high tolerability of ATO in healthy tissues (illustrated by its current clinical use), is counterintuitive and intriguing, as many healthy tissues tend to rely on OXPHOS to produce ATP. This targeted inhibition raises the question of how microenvironmental effects would modulate the efficacy of ATO in solid tumors, and whether the presence of gradients would generate spatially resolvable populations resistant to its metabolic reprogramming.

To address this, we use a 3D microphysiological system (MPS) consisting of an HGSOC spheroid embedded within a collagen hydrogel adjacent to a media-exchange channel (and, where appropriate, an endothelial vessel) in a microfluidic device. This system recapitulates tumor metabolic heterogeneity and enables longitudinal, readouts of ECM invasion, viability, and metabolic state. Using this system, we assessed spatial differences in spheroid response to ATO and evaluated metabolic reprogramming through different readouts (cell viability, proliferation, migration, semi-quantitative metabolic imaging, and mitochondrial network reorganization).

Our results revealed that ATO cell death and metabolic reprogramming in HGSOC spheroids. Specifically, ATO was effective at reducing invasion area, invasion distance and cancer cell proliferation in spheroids in our model. Further, we observed that ATO switched cell migration modes from individual to collective, and over time disrupted mitochondrial networks in the spheroid invasive fronts. Finally, our results show that ATO was able to decrease levels of YAP in the spheroid edge alone. This result provides a potential mechanism behind the decrease in spheroid invasion and opens the door to further studies on harnessing OXPHOS inhibition as potential maintenance therapy against HGSOC.

## Results and Discussion

### Atovaquone, an OXPHOSi, decreases viability and invasion of HGSOC spheroids but does not damage endothelium

Tumors display an array of metabolic phenotypes and specifically display different levels of reliance on oxidative phosphorylation (OXPHOS) according to tumor cell needs and access to nutrients^24^. Here, we wanted to explore if metabolic reprogramming of tumors could be leveraged to target solid tumor invasion and dissemination. To this end we used cell lines traditionally used to model high-grade serous ovarian cancer (HGSOC), and OXPHOS inhibitor Atovaquone (ATO), given the recent focus of these kinds of therapeutics as a potential maintenance therapy for HGSOC and other cancers^25–27^. Although most OXPHOS inhibitors have not proven useful as anti-cancer therapeutics due to their systemic toxicity, ATO (an FDA-approved complex III inhibitor of the mitochondrial transport chain typically used to treat malaria) does not carry that toxicity and therefore has shown promise as an anti-cancer agent in conventional 2D *in vitro* models^25^. The premise behind its anti-tumor activity is that tumor cells require OXPHOS^28^, and it is hypothesized that they are more dependent on ATP produced by this metabolic process than normal cells^29^. Therefore, theoretically ATO at the right dosage could be used to selectively target tumor cells. However, the mechanisms behind its anti-tumor activity are not well understood and largely studied in 2D only^25, 27^. Nutrient and oxygen access in 3D solid tumor environments are known to affect proliferation, migration mechanisms and other cell functions, which could impact ATO effectiveness against solid tumors^30^. Furthermore, in 3D solid tumor environments not all cells have the same metabolic needs and therefore display different metabolic phenotypes, which could also impact OXPHOS reliance, and therefore decrease ATO effectiveness^31^.

To better understand the effectiveness of ATO in a system more closely recapitulating the metabolic heterogeneity of tissues than 2D cell cultures, we generated a microphysiological system (MPS) to mimic the solid tumor heterogeneity of HGSOC and investigate tumor response to ATO and healthy tissues that are highly dependent on OXPHOS. Our model consists of an HGSOC tumor spheroid embedded in a collagen hydrogel within a microfluidic device, and a surrogate blood vessel surrounded by a collagen hydrogel that mimics the extracellular matrix (ECM) (Fig. 1A, center). Our choice to use tumor spheroids was due to their inherent self-generated biological gradients (e.g., lactate, oxygen, and nutrients) that lead to a variety of cell populations at diverse metabolic states (i.e., proliferative, quiescent, necrotic) despite their clonal origin, more closely resembling those of solid tumors (Fig. 1B). The metabolic phenotype of the cells will also dictate energy available for high energy-demanding cell processes (i.e. migration and proliferation). For these experiments, we used established cell lines that have traditionally been used as models for HGSOC *in vitro* (i.e., OVCAR5 and SKOV-3). The reasons behind this choice are two-fold: 1) Among the cell lines to mimic HGSOC, these display a higher variety of metabolic phenotypes and invasive behaviors than other options (e.g., OVCAR3, PEO1), 2) literature available for ATO has often used these two cell lines, which allows us to compare our results to those published. Altogether, the use of a spheroid allows us to capture more biological processes that are potentially impacted by metabolic changes and ultimately understand how ATO may impact the diverse metabolic phenotypes present in the TME. Spheroid cell numbers were chosen to produce a significant necrotic core to ensure we recapitulated a variety of metabolic phenotypes in the same system.

**Figure 1:**
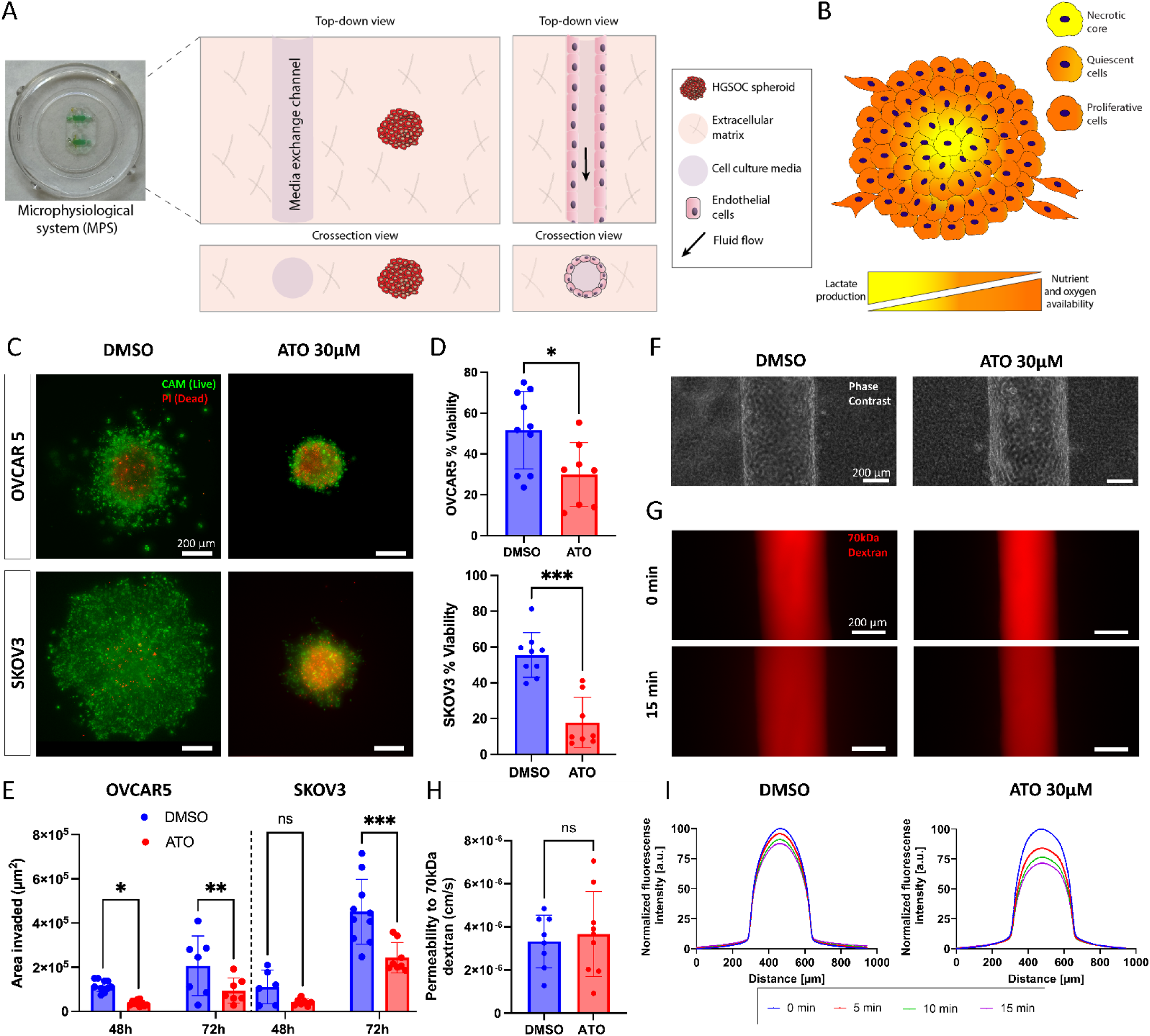
OXPHOSi ATO reduced cell viability in tumor spheroids but not in healthy blood vessel models. **(A)** Photography of microphysiological device (left) and schematic illustration (center) of MPS used to test the effects of an OXPHOSi, atovaquone (ATO) 30µM, on HGSOC spheroids. This model contains an HGSOC spheroid embedded in a collagen hydrogel within the chamber of the MPS, and a tubular structure for media exchange. A schematic illustration (right) of the MPS used to model an endothelial cell vessel in 3D. The chamber contains a collagen hydrogel containing an empty tubular channel lined with HUVECs. Media containing ATO is then perfused through the vessel. **(B)** Schematic illustrating the biological gradients and the multiple cell populations of a tumor spheroid. Cells in the periphery of the tumor spheroid have greater access to biological gradients, driving invasion and proliferation. Meanwhile, cells in the center represent a necrotic core due to limited access to nutrients and oxygen. **(C)** Fluorescent images of OVCAR5 and SKOV-3 spheroids stained with calcein AM (CAM) and propidium iodide (PI) to determine viability after treatment with ATO 30 µM **(D)** Quantification of % viability in OVCAR5 and SKOV-3 spheroids treated with ATO compared to control. **(E)** Quantification of area invaded by spheroids (excluding spheroid body) at 48 and 72h. **(F)** Phase contrast images of endothelial cell-engineered vessels treated with ATO vs control. **(G)** Representative fluorescent images of biomimetic blood vessels at 0 min and 15 min perfused with TexasRed-70 kDa dextran to determine if ATO affects the permeability of the vessel. **(H)** Plot profile of the dextran fluorescence intensity. **(I)** Permeability coefficient calculated from dextran diffusion data. Unpaired t-test was used for (D) and (I), One-way ANOVA was used for (G) n<3 for all experiments, *p < 0.05; **p < 0.01; ***p < 0.001.

We also chose HUVEC since they are primary endothelial cells commonly used to mimic the endothelial barrier in microfluidic models^32^ and can serve as a healthy cell control with high sensitivity to ROS derived from OXPHOS decoupling^33^. Finally, the presence of a naturally derived ECM allows us to better capture cell-ECM dependent interactions that regulate important processes in tumor progression, namely, mechanotransduction, cell adhesion, proliferation, and migration^34^.

As a first approach to determine if ATO has effects on HGSOC spheroids, we treated SKOV3 and OVCAR5 spheroids with ATO at the IC50 calculated in prior 2D *in vitro* studies (30 μM)^35^. In those *in vitro* and *in vivo* HGSOC models, ATO decreased the proliferation and viability of HGSOC cells^25^. Live-dead staining revealed that ATO-treated OVCAR5 and SKOV-3 spheroids showed more cell death compared to the vehicle control (Fig. 1C). Through area quantification, ATO-treated OVCAR5 spheroid viability decreased from 51.67 ± 18.93% to 29.91 ± 15.7% (*p=0.02), and for SKOV-3 from 55.55 ± 12.51% to 17.82 ± 14.1% (***p<0.01) (Fig. 1D). Notably, only the cells at the very edge of the SKOV-3 spheroids remained alive, whereas cells invading the ECM were viable yet fewer than in the control. Spheroid viability was not halved upon ATO treatment at IC50 dosages, underscoring the value of testing anti-cancer agent cytotoxicity in 3D systems. The differences in sensitivity for each of the spheroid types (42% decrease in OVCAR5 and 68% decrease in SKOV3) are consistent with the reported differences in OXPHOS reliance for these cell types^36^. Spatially, we also observed that the dead cells are mainly localized in the center of the spheroid, whereas cells invading the surrounding ECM were decreased, yet remained alive (Fig. 1E). In theory, one would expect that cells with higher access to oxygen to have more OXPHOS reliance (i.e., more external to the spheroid) and therefore show more sensitivity to treatment than cells closer to the core. However, our results could be explained by the lower strength of ATO as an OXPHOS inhibitor^37^. Given the only partial inhibition of OXPHOS by ATO, cells located at the outside may have enough energy-producing pathways still available to them to stay viable, whereas cells located more inward in the spheroid are likely more energy-depleted and therefore more susceptible to the effects of ATO.

Following the same logic, we hypothesized that ATO would be effective at reducing invasion for both cell lines, since migratory cells would all have a more OXPHOS-dependent phenotype. Indeed, for OVCAR5 we observed an increase in ECM invasion by migratory cells that was slower than that of SKOV-3 (1.77-fold increase in area invaded by cells from 48-72h compared to a 4.05-fold increase respectively) (**p=0.005). Treatment with ATO hindered this invasive potential similarly for both cell types (54% and 46% reduction of invaded area) (p=0.7) respectively, indicating that ATO is consistently effective at reducing cell invasion (Fig. 1G). This prompted the question of whether an OXPHOS-reliant healthy cell type would show increased toxicity to ATO in our system.

An important pitfall of OXPHOS inhibitors in clinical trials was the systemic cytotoxicity they caused as they inhibited OXPHOS in healthy cells that rely upon this metabolic pathway for survival^22, 38^. Even though ATO is clinically approved and commonly used for malaria, the concentrations used for its anti-cancer properties differ from its anti-malarial applications. Thus, we wondered whether the concentrations used for anti-cancer applications would result in toxicity for non-malignant cells that heavily rely on OXPHOS, such as endothelial cells. Blood vessels are highly OXPHOS dependent during angiogenesis, display a clear response to excess reactive oxygen species (ROS) and will be exposed to ATO during drug delivery to tissues, making them an interesting case study for testing potential toxicity of ATO in a healthy tissue. Therefore, we used our highly established biomimetic endothelial cell vessel model (Fig. 1A, right), with HUVECs and tested the effects of ATO on vessel integrity, barrier function and cell-cell junctions. In this model, if ATO were toxic to the endothelial cells, we would expect floating dead cells and gaps in the vessel as they contract, decrease cell-cell junctions, and potentially detach from the collagen tubular channel. However, no floating cells or major gaps in the vessels were seen (Fig. 1F) compared to the DMSO control. To determine potential functional impacts on the barrier function for our surrogate blood vessel, we performed a dextran diffusion test using fluorescently labeled 70 kDa dextran (equivalent to BSA in hydrodynamic radius). This test revealed no major leakage in ATO-treated vessels compared to the control, indicating the vessel barrier function is not affected (Fig. 1G, 1H). Quantification of permeability coefficient between ATO-treated vessels and control showed comparable values for both: for ATO-treated vessels, permeability was 3.67·10^−6^ ± 1.97·10^−6^ cm/s, and for vehicle control (DMSO) it was 3.32·10^−6^ ± 1.23·10^−6^ cm/s (p=0.67) (Fig 1I). We also looked at vessel confluence as another measurement of viability using immunofluorescence for F-actin, and we did not see any major gaps in the ATO-treated or control vessels visualized by F-actin (Fig. S1), confirming the integrity of the vessels. Overall, this data indicates that at concentrations relevant for targeting cancer cells, ATO does not impact the integrity and function of normal endothelial cell vessels.

### HGSOC spheroids undergo metabolic reprogramming upon OXPHOS inhibition with ATO

Next, we wanted to confirm that ATO produces a functional and significant metabolic shift of the HGSOC spheroids in its mechanism of action, consistent with our hypothesis. During OXPHOS, cells oxidize NADH and FADH_2_ to produce ATP, releasing NAD and FAD in this process. Conversely, cells with a prevalent glycolytic phenotype will reduce NAD^+^ to produce ATP, releasing NADH in this process (Fig. 2A). To semi-quantitatively determine the metabolic phenotype of the HGSOC spheroids treated with ATO, we used metabolic imaging, a technique that captures the autofluorescence of both NADH and NADPH in one wavelength (hereafter referred to as NAD(P)H for simplicity), and FAD in another wavelength, which is quantified to determine the optical redox ratio (ORR) of the spheroids ^39^. Here, we used a modality of metabolic imaging that only requires a conventional fluorescence microscope to produce a measurement of spheroid FAD and NAD(P)H autofluorescence for the entire spheroid, and therefore a metabolic profile for the spheroid body, with the tradeoff that signal for single cells is too low to be detected in this setup ^40^. Autofluorescence data is then quantified to calculate ORR as described in materials and methods. ORR is interpreted as follows: an increase in ORR suggests an increase in NAD(P)H. Increase in ORR is consistent with an increase in the NAD(P)H/FAD ratio (Fig. 2B).

**Figure 2:**
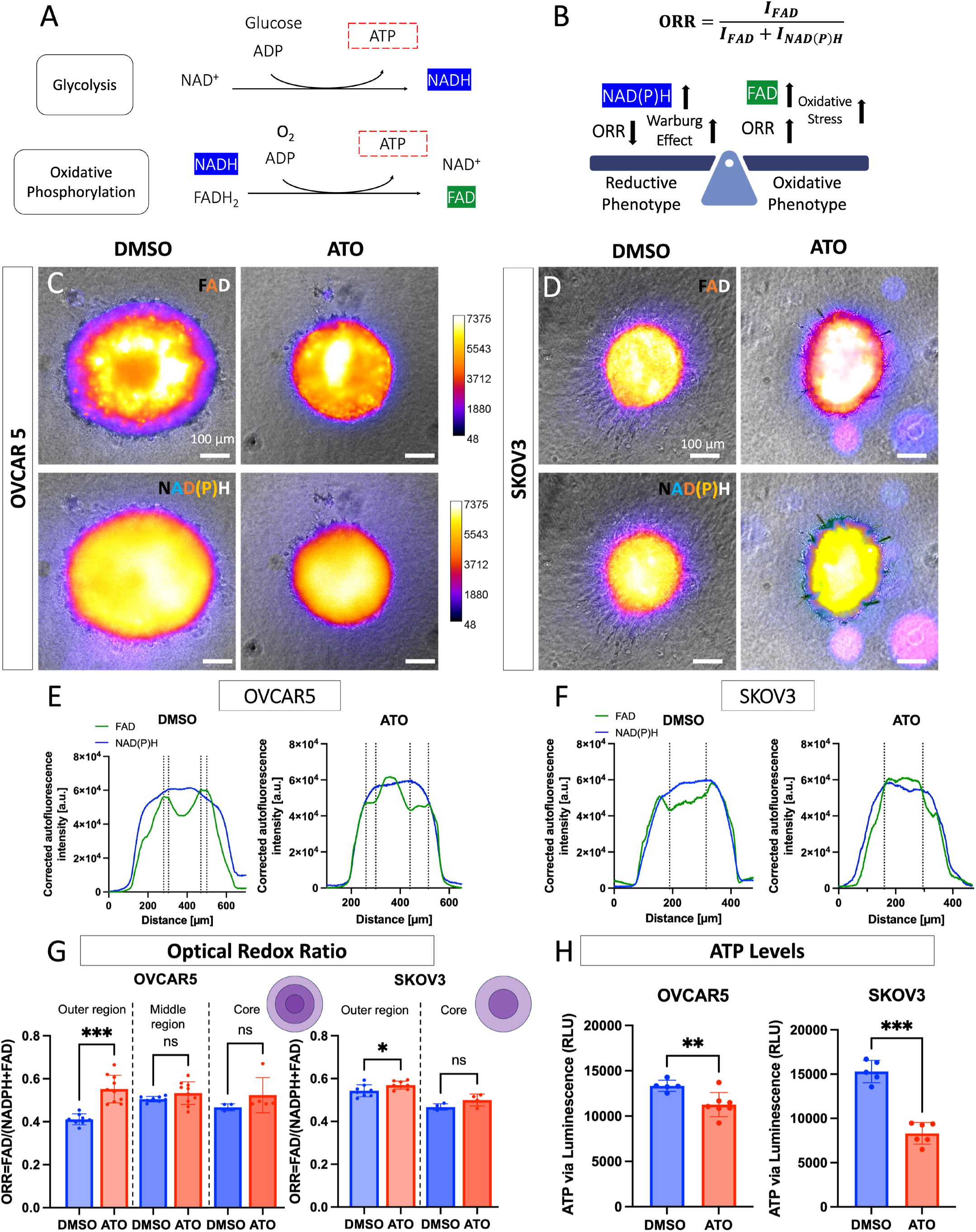
OXPHOSi ATO induced metabolic reprogramming in tumor spheroids. **(A)** Schematic of consumption and production of metabolic species highlighted by their autofluorescence, including production of adenosine triphosphate (ATP) in glycolysis and oxidative phosphorylation (OXPHOS) metabolisms. **(B)** Illustration of optical redox ratio (ORR), flavin adenine dinucleotide (FAD), and nicotinamide adenine dinucleotide phosphate (NAD(P)H) levels according to the activity of the prevalent metabolic phenotype. **(C-D)** Representative images of OVCAR5 and SKOV3 spheroids capturing the autofluorescence of NAD(P)H and FAD species after 48h of treatment with ATO and DMSO. Dotted lines represent the different areas of the spheroids. **(E-F)** Plot profiles of corrected autofluorescence intensity of FAD and NAD(P)H for OVCAR5 and SKOV3 for DMSO and ATO treatments. **(G)** ORR values for OVCAR5 and SKOV3 spheroids for ATO and DMSO treatment groups. **(H)** Levels of ATP via a luminescent reporter of OVCAR5 and SKOV3 spheroids after 48h of ATO treatment determined by a luminescent reporter. Welch’s t-test was used for (E), while the unpaired t-test was used for (F). *p < 0.05; **p < 0.01; ***p < 0.001. Except for SKOV3 ORR (n=3), n=>5.

We initially calculated global ORR treating whole spheroids as a single entity (Fig S2) by generating an ROI that was the same size and shape as the spheroid body, we saw that ORR significantly increased for OVCAR5 (0.12 ± 0.07 for control vs 0.15 ± 0.03 for ATO) but surprisingly decreased for SKOV3 (0.45 ± 0.06 for control vs 0.21 ± 0.03 for ATO-treated). Conversely, SKOV3 increased ORR upon treatment, therefore seeming to shift its metabolism more toward OXPHOS. When we inspected the images in more detail (Fig. 2C), we observed that the signal, especially for FAD, was not constant throughout the spheroid, and that several regions consistently appeared across different spheroids of the same cell type and treatment condition. Therefore, we hypothesized that changes in autofluorescence could reveal different metabolic phenotypes across the spheroids and indicate where ATO would be more effective.

Untreated OVCAR5 spheroids displayed a central region defined by an FAD dip and NAD(P)H signal was high, an intermediate region where FAD peaked and produced a small plateau, before quickly decreasing again in the outer region (Fig 3E left). Conversely, NAD(P)H produced a dome-shaped curve. Upon treatment, FAD lost its rounded M-shaped curve and instead peaked in the central region (Fig 3E right). The regions showcasing this increase in FAD coincide with the propidium iodide signal in our viability studies. In SKOV3 spheroids, plot profiles of FAD and NAD(P)H were analogous to those of OVCAR5. However, an intermediate region was not apparent. We consistently observed that while the FAD curve had a smaller area under the curve in untreated spheroids, upon ATO treatment areas were much more comparable, and area under the FAD curve was larger in the outer regions of the spheroid. These results suggest that there is a distinct metabolic phenotype in an intermediate region of OVCAR5 spheroids that does not appear in SKOV3 spheroids. Our data suggest that this region undergoes changes in autofluorescence, yet no significant changes in metabolic phenotype are observed, which could suggest a metabolic switch away from these main pathways.

**Figure 3:**
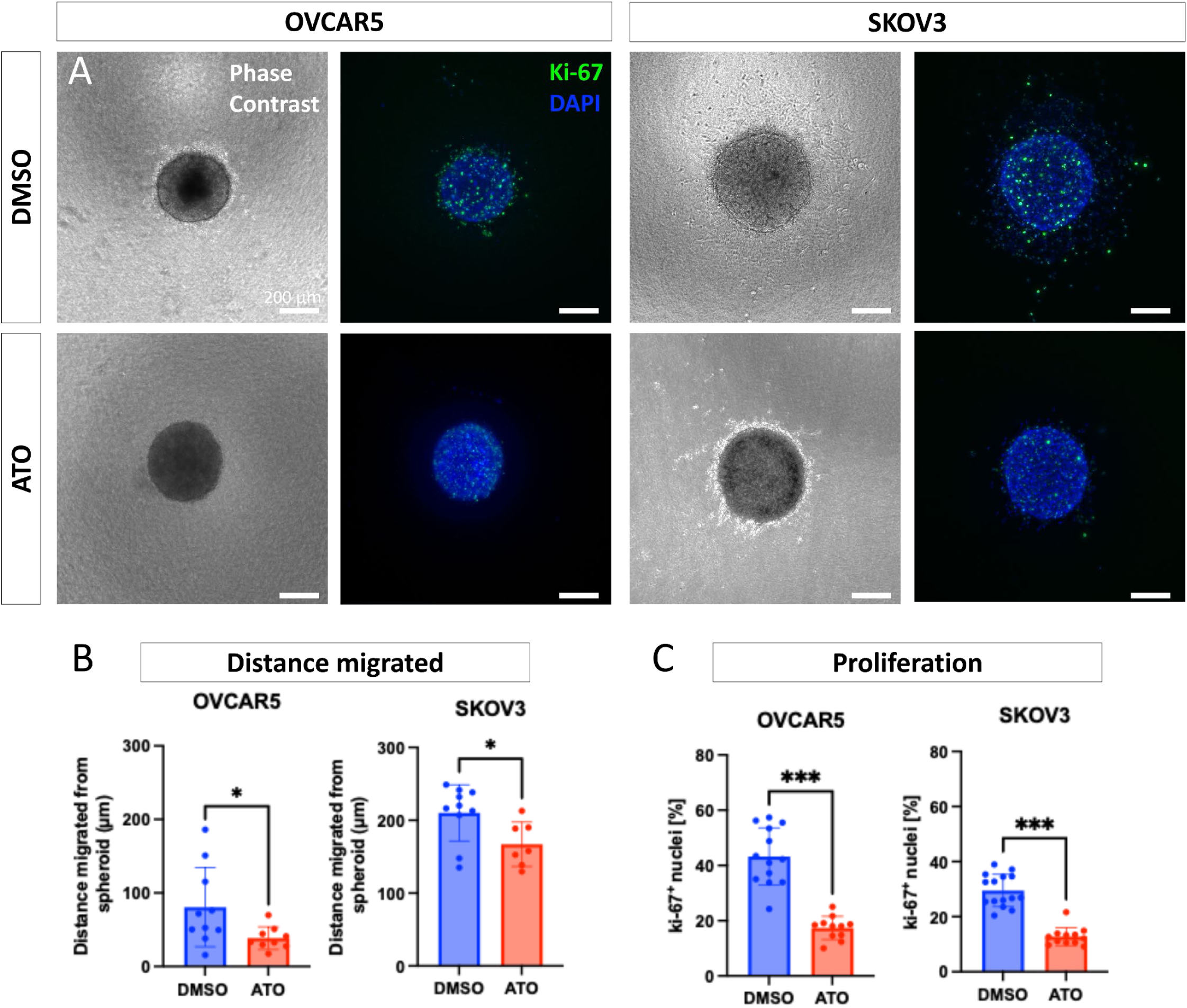
ATO decreases migration and proliferation of HGSOC spheroids. **(A)** Representative phase contrast and fluorescent images of ki67 (a proliferation marker) for OVCAR5 and SKOV3 spheroids at 48h for ATO and DMSO treatment groups, where different phenotypes and migration patterns can be observed. **(B)** Quantification of distance migrated by the cells in the periphery of the spheroids into the ECM. **(C)** Quantification of the Ki-67 signal in the HGSOC spheroids for ATO and DMSO. Welch’s t test was used for (B) and (C). *p<0.05; p***<0.001; n=3 spheroids.

We calculated ORR for each region using the area under the curves and observed that the ORR in the outer region consistently increased significantly in both spheroid types upon treatment (for SKOV3 0.54 ± 0.028 vs 0.57 ± 0.019, *p=0.04) (0.41 ± 0.025 vs 0.55 ± 0.064, ***p<0.001). This increase indicates an accumulation of FAD in the region. FADH_2_ synthesis would require energy from the mitochondrial transport chain. Instead, cellular stress from ATO transport chain results in ROS production and suggests a global shift away from an OXPHOS-reliant phenotype in this region. To support the metabolic rewiring away from OXPHOS, we hypothesized that ATP production would be significantly decreased. Consistent with the prior data, CellTiterGlo (which directly measures ATP) revealed a greater difference in the levels of ATP between the ATO and control groups for the SKOV3 spheroids (ATO: 8.31 ± 1.22 ·10^3^, DMSO:15.29 ± 1.27·10^3^, 45.6% decrease, ***p<0.001) compared to the OVCAR5 spheroids (ATO: 11.27 ± 1.34·10^3^, DMSO: 13.34 ± 0.62·10^3^, 15.5% decrease, ***p<0.001). Altogether, ATO appears to induce a metabolic shift in HGSOC spheroids, altering metabolic phenotypes and decreasing energy production.

Interestingly, SKOV3 images showcase how ATO crystals get formed within the hydrogel and around the spheroid. We have observed these in absence of cells only in ATO-treated conditions (Fig S3), due to the low solubility of ATO in aqueous media. Decreased NAD(P)H autofluorescence can be observed around the areas around these crystals, indicating that ATO has a direct impact on these species.

### Invasive and proliferative behavior of HGSOC spheroids is inhibited with ATO

Altered tissue metabolisms are defining features of cancer, that impact both proliferation and migration^4, 16^. However, it has been suggested that cells can only perform one high energy-demanding process at a time, (i.e., “go or grow”) namely migration or proliferation^39^. Both processes, in fact, are highly affected by a 3D microenvironment^40^ – cells in 3D proliferate and migrate less than in 2D^13^. In the case of proliferation, it is highly impacted by substrate stiffness and therefore lowered in 3D microenvironments compared to highly stiff plastic petri dishes^41^. This behavior can impact the effectiveness of many chemotherapeutics, which mainly target proliferating cells, and drives chemoresistance specifically in ovarian cancer^42^. On the other hand, cancer cell migration in 3D is a process that is highly energy demanding^43^, due to the need for cells to constantly rearrange their cytoskeleton and secrete metalloproteinases that will allow them to overcome extracellular matrix obstacles^44^. There are stark differences in ATP consumption between individual cells migrating and leader cell migration, where the latter are less affected by energy fluctuations. While it is known that an unfavorable glycolytic switch is not only beneficial but necessary for cancer cell motility^16, 43^, the glycolysis-OXPHOS balance regulation in leader cells is unclear^40^. Long-distance cell migration is a high energy-demanding process^16^, since it requires deep bursts of energy on demand to overcome environmental constraints. These energy bursts may be achieved by having a high energetic plasticity^7^, defined as the ability for cells to dynamically switch from an oxidative phenotype to a highly glycolytic one^43^. We specifically posit that the maximum distance migrated by individual invasive cancer cells will be significantly affected by ATO. Therefore, our MPS will more closely inform of the effectiveness of ATO in solid tumors and could reveal spatial patterns of ATO effects on solid tumors.

Therefore, we first measured the 6 longest migration distances of cells outward from the edge of the spheroid body and averaged them to calculate the maximum migrated distance for each spheroid type daily until 72h of culture. Distance quantifications showcased increased migrations over time, with SKOV3 spheroids displaying a more migratory phenotype than OVCAR5 cells, representing heterogeneous phenotypes of HSGOC and consistent with prior reports^45^. Indeed, we observed that both migratory phenotypes were hindered in ATO-treated conditions. Treatment affected maximum migrated distance for both OVCAR5 and SKOV3 (80.71 ± 53.81 µm vs 38.55 ± 15.37 µm and 210.0 ± 38.66 vs 167.2 ± 30.51 µm, respectively). Interestingly, longest migration distances were only reduced by about 40 µm in both phenotypes, indicating that ATO has a moderate effect on metabolic plasticity, yet we did note fewer cells undergoing individual cell migration in the treated conditions.

Next, we assessed whether ATO effectively targets proliferating cells, which should in theory be more abundant at the periphery of the spheroid body and not as abundant away from the spheroid body. (Fig. 3D, E) Staining of ki67, a proliferation marker, revealed that the largest concentration of proliferating cells was located on the edge of the spheroid body but not among invasive cells in the surrounding ECM. OVCAR5 displayed a higher percentage of proliferation compared to SKOV3 cells and ATO effectively decreased the percentage of cells positive for ki67 in both spheroid types. Specifically, the percentage of ki67 positive cells in OVCAR5 decreased 60% (from 43.2 ± 10.4% to 17.4 ± 4.20%), and in SKOV3 spheroids the decrease in ki67+ % was a comparable 57% (from 29.5 ± 5.87% to 12.8 ± 3.30%). These results showcase that ATO is moderately effective at reducing metabolic plasticity in solid tumors, resulting in a lower number of low-distance migratory cells; but it is especially effective at reducing cell proliferation, both in the body of the spheroid and for cells that have invaded the surrounding ECM.

### Atovaquone disturbs mitochondrial networks of HGSOC spheroid invasive fronts

Mitochondria play an especially important role in highly energy-consuming cellular processes like cell migration. These highly dynamic organelles are organized in networks, which are dynamic and interconnected webs that remodel in response to cell demands^46^. Migrating cells need to adjust their bioenergetics to navigate the structural and mechanical cues of their surrounding ECM. To migrate within a 3D microenvironment, cells must support the high energetic expenditure of cytoskeletal reorganization and force generation^8, 16, 47^. During cell motility, mitochondria localize in the lamellipodia of the cells, creating elongated networks that allow them to efficiently produce enough ATP via OXPHOS to meet the locally high bioenergetic demands. Although the role of metabolism in cancer cell motility is still emerging^40^, evidence suggests that cell bioenergetics will shift according to the type of cell migration: either individually or collectively as multicellular groups^48^. During collective migration, the leader cell consumes more energy to overcome the resistance in front of it than follower cells. Follower cells can then take over as leaders to support collective migration while minimizing total energy expenditure^40^. This dynamic makes collective migration a less costly process than individual cell migration^49^.

Following our finding that ATO decreased ATP levels as well as decreased migration in SKOV3 spheroids predominantly, we were interested in the effects of OXPHOS inhibition on the distribution and morphology of mitochondrial networks. Here, we hypothesize that ATO treatment will disturb the mitochondrial networks within the cells and affect their capability to migrate individually in the short term. At longer time points, we expect that the mitochondria will appear more fragmented with treatment and will be localized around the nucleus due to the stress response caused by the inhibition of OXPHOS. To observe the effects of ATO in migratory cells, we inspected the mitochondrial networks in migratory fronts sprouting from the spheroid. To this end, we used Mitotracker staining to investigate the organization of mitochondrial networks over time and used a non-toxic live cell staining as an individual cell indicator. This staining combination allowed us to determine the organization and local distribution of the mitochondria at a single-cell level at different timepoints. A critical mass of cell migration outside of the spheroid was only accessible for imaging in the SKOV3 spheroids due to their highly migratory phenotype.

We imaged the samples at 48 h and 72 h to track the changes of migration, cell morphology, and mitochondrial networks. In the control SKOV3 spheroids at 48h, we observed predominantly elongated mitochondrial networks polarized on the opposite sides of cell nuclei, a critical step for individual cell migration^50^. At the same time point, the cells treated with ATO are more rounded and closer together, consistent with a more epithelial phenotype and a pattern of collective migration. Cells with polarized mitochondria, such as the ones observed in DMSO control, were less abundant. To quantify these cell phenotypes across multiple images, we used the Mitotracker signal to calculate the cell aspect ratio. Cells treated with ATO showed a significantly lower aspect ratio (2.2 ± 0.60) compared to the vehicle control (5.5 ± 2.2), supporting our claim that the mitochondria are not as localized in the invasive front of the cells, and that the migration patterns have been shifted based on the bioenergetic restrictions imposed by ATO treatment.

At 72h, we see a complete disorganization of mitochondrial networks in the cells treated with ATO compared to the control. In the control condition we observed the persistence of the mitochondrial polarization pattern we described at 48h. Additionally, mitochondria in control conditions appear to constitute a more cohesive network, whereas in the ATO treated condition they have a more spread-out punctate appearance, consistent with prior reports of cell stress and apoptotic phenotypes.

Although we can only infer the bioenergetic profile of cells from Mitotracker and nuclei data, the field has indicated that metabolic flexibility allows cells with high metabolic demands (e.g., cells undergoing individual cell migration) to switch their metabolism from OXPHOS to glycolysis to overcome the constraints of 3D cell migration. ATO treatment would hinder this metabolic flexibility, impinging more on cells undergoing individual cell migration, and potentially less those in collective cell migration^51^. Our data suggests that the metabolic restriction of ATO is enough to hinder cell migration in both cell types, and that the mitochondrial stress experienced by those cells may drive them into apoptosis^52^, therefore effectively hindering cell migration.

### OXPHOSi affects expression of YAP1 in the cells towards the outer layer of HGSOC spheroids

The Hippo pathway is a central regulator of cell proliferation, survival, and invasion that is frequently dysregulated in cancer^53,54^. Its activity is closely linked to cellular metabolism and mitochondrial function^55^. Under energy-rich conditions, Hippo signaling pathway is inactivated, allowing its transcriptional co-activator YAP1 to accumulate in the nucleus and promote the expression of genes associated with tumor growth and progression. Conversely, when ATP production is impaired, metabolic sensors can inhibit YAP1 activity either directly or through activation of the Hippo pathway, resulting in YAP1 phosphorylation, cytoplasmic sequestration, and degradation^56,57^. Since we demonstrated that atovaquone disrupts mitochondrial respiration and reduces cell proliferation and migration, both processes that are highly dependent on ATP production, we hypothesized that part of its antitumor activity may be mediated through modulation of YAP1 nuclear localization and total protein levels.

To investigate this possibility, we evaluated the spatial distribution of total YAP1 (t-YAP1) protein in OVCAR-5 spheroids following atovaquone treatment using immunofluorescence imaging. In control spheroids, t-YAP1 expression was enriched at the spheroid periphery, corresponding to the highly proliferative and invasive tumor edge, as described in other studies^58^. In contrast, ATO-treated spheroids exhibited reduced peripheral t-YAP1 intensity. DAPI and phalloidin signals were more abundant within the spheroid core, consistent with higher cellular packing density in this region (Fig. 5A-B). To normalize t-YAP1 expression to cell density, a horizontal slab passing through the center of each spheroid cross-section was selected, and fluorescence intensity profiles for stainings were generated using ImageJ to quantify and compare signal distribution across the spheroid diameter (Fig. 5C), therefore accounting for intensity changes in spherical shapes. DAPI fluorescence was used to normalize the t-YAP1 signal. The YAP1/DAPI ratios at the spheroid edge, corresponding to the regions displaying the highest YAP1 signal intensity, were then compared between control and ATO-treated spheroids. This analysis revealed a significant reduction (∼40%) in relative t-YAP1 signal intensity at the spheroid periphery following atovaquone treatment (Fig. 5D). To ensure that DAPI intensity was consistent throughout treatment groups, DAPI signal was further normalized to phalloidin intensity. No significant differences were observed between control and atovaquone-treated spheroids, supporting the validity of the normalization approach. Together, these findings indicate that atovaquone reduces t-YAP1 levels at the tumor edge in HGSOC spheroids, a region characterized by active proliferation and invasion.

The mechanism underlying this spatially restricted reduction in t-YAP1 remains to be fully elucidated. Based on our findings and the current literature, two main hypotheses can be proposed to explain the reduction in YAP1 levels observed following atovaquone treatment. One is that ATO-induced ATP depletion activates AMPK, which may suppress YAP1 activity and promote its degradation either directly or through Hippo pathway activation^56^. A second possibility is the modulation of YAP1 levels by oxidative stress. Oxidative stress has been reported to activate the Hippo signaling pathway through multiple mechanisms, influencing its core regulatory components and promoting the cytoplasmic sequestration and degradation of YAP1^59^. Since the ORR is higher at the spheroid periphery, the observed reduction in YAP1 levels may be associated with a more oxidative microenvironment in this region following atovaquone treatment. We attempted to further characterize the downstream consequences of reduced YAP1 signaling. However, the limited availability and specificity of antibodies suitable for immunofluorescence restricted our ability to comprehensively assess YAP1 downstream targets. Moreover, bulk analytical approaches such as western blotting and RT-qPCR failed to detect significant changes in YAP1 expression or activity at the whole-spheroid level (Supplemental Fig.4), likely because the ATO-induced decrease in t-YAP1 was primarily confined to the spheroid periphery and therefore diluted in bulk lysates. These observations underscore the importance of spatially resolved methods for studying YAP1 regulation, whose activity appears to depend not only on intrinsic cellular signaling but also on the local tumor microenvironment and the heterogeneous architecture of the tumor. Future studies employing spatial transcriptomics or region-specific protein extraction may help delineate the full mechanistic consequences of atovaquone-mediated YAP1 suppression at the tumor edge. Collectively, our results support the use of physiologically relevant three-dimensional models to better understand how metabolic therapies influence YAP/TAZ signaling and tumor behavior.

**Figure 4:**
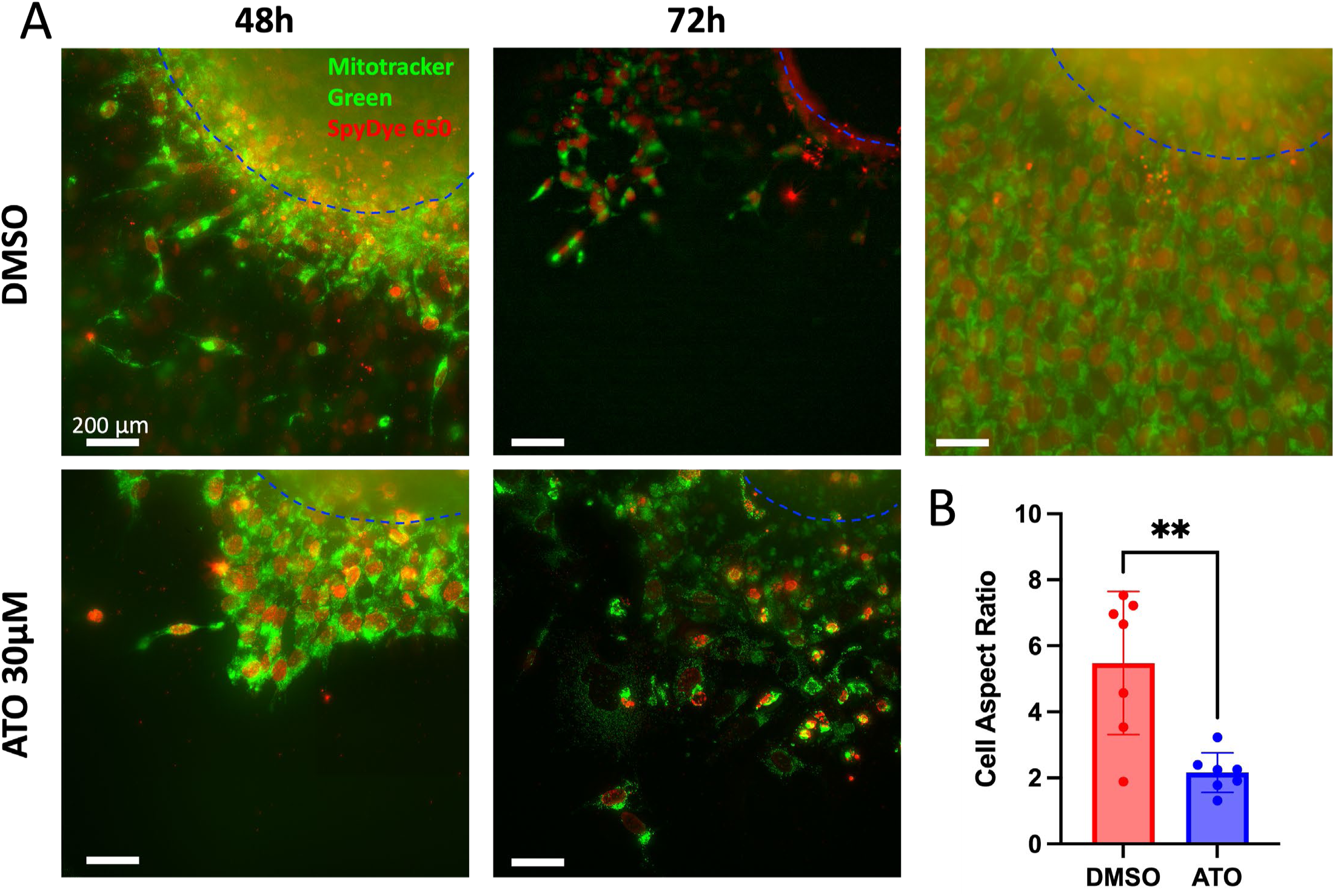
Atovaquone disrupts mitochondrial networks in SKOV3 spheroids. **A)** Representative fluorescent images of cells in the periphery of SKOV3 spheroids treated with ATO and DMSO at 48h and 72h stained with Mitotracker Green (green) to visualize the mitochondrial networks and SpyDye 650 (red) as a nuclear counterstain. The top right image of the spheroid at 72h was taken at a different focal plane than the one in the middle, depicting zonal differences in migration. **B)** Quantification of cell aspect ratio of migrating cells at 48h. Spheroid body was marked with a dashed blue line. Welch’s t test was used. **p=0.006, n=3.

**Figure 5:**
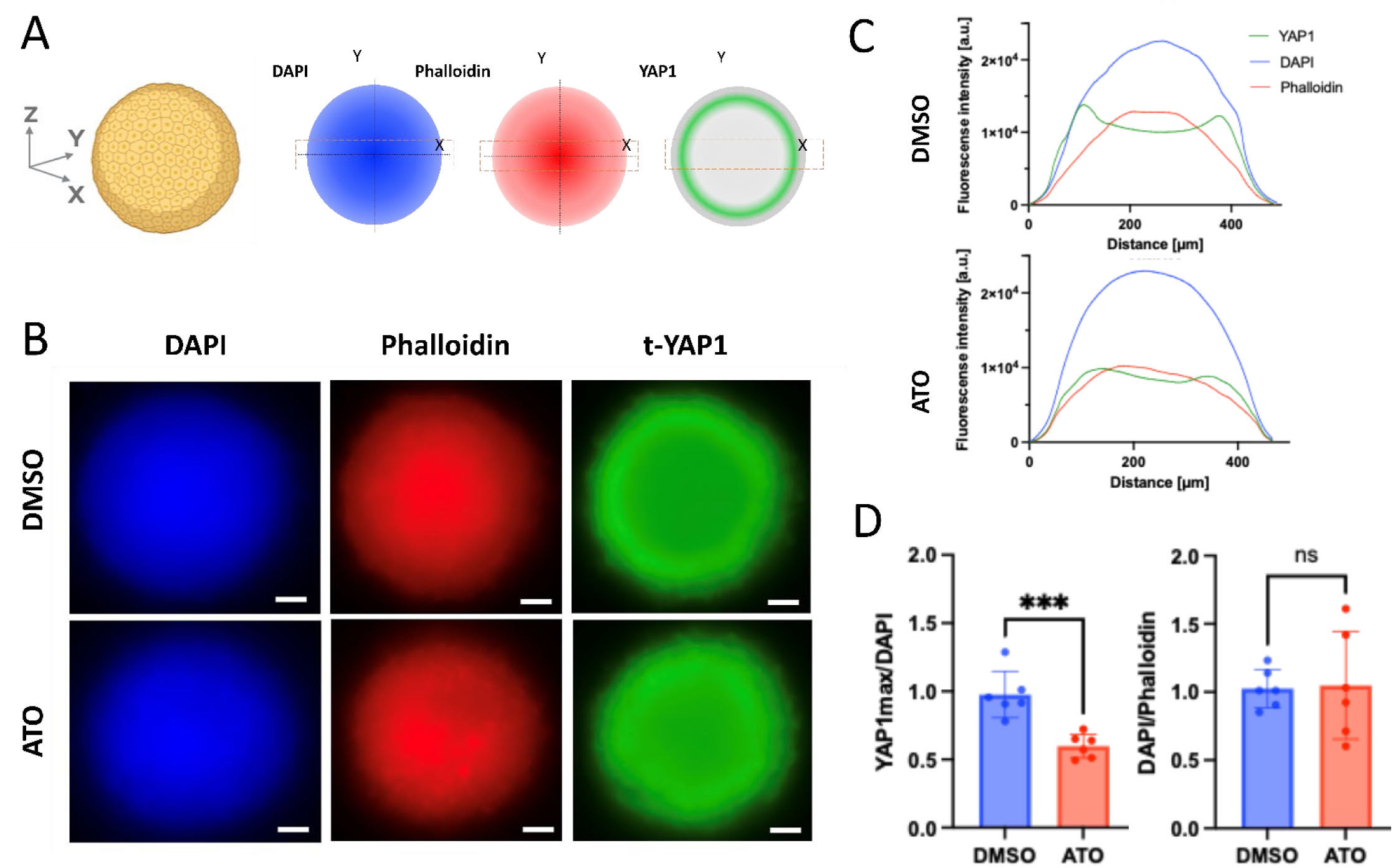
Atovaquone decreases YAP1 expression at the edge of HGSOC spheroids. A) Illustration of the signal intensity across a spheroid cross section for each immunofluorescence staining. The dashed rectangle indicates the regions where the YAP1 signal reaches maximal intensity and corresponds to the site used for ratio quantification shown in (D). B) Representative images of YAP1 staining together with DAPI and phalloidin (actin) staining. C) Plot profiles of fluorescence intensity of YAP1, DAPI, and phalloidin for spheroids treated with DMSO and ATO. D) Quantification of the YAP_max_/DAPI and DAPI/Phalloidin signal ratio. Each point in the graph represents a single ratio. p***<0.001, *n* = 3 spheroids per treatment group. Scale bar = 200 µm

## Conclusion

Here, we developed a microphysiological system (MPS) that enables a reductionist yet physiologically relevant recapitulation of the heterogeneous cell states and bioenergetic landscapes characteristic of metastatic HGSOC solid tumors. As a proof of concept, we selected a metabolic inhibitor expected to differentially affect distinct metabolic subpopulations within this system. Atovaquone (ATO), an FDA-approved antimalarial drug and inhibitor of oxidative phosphorylation (OXPHOS), has recently been repurposed for its potential anticancer activity. Notably, compared to other OXPHOS inhibitors, ATO demonstrates relatively low toxicity in healthy tissues while still effectively impairing mitochondrial respiration.

Using this platform, we evaluated the effects of ATO on HGSOC spheroids at near single-cell spatial resolution within a 3D collagen-rich microenvironment that mimics tumor progression in the ovary. Our results show that ATO does not uniformly suppress tumor viability; rather, it selectively constrains metabolically vulnerable regions, particularly the proliferative and invasive spheroid periphery. These regions are critical drivers of tumor progression, underscoring the importance of spatially resolved analysis in evaluating metabolic therapies.

Specifically, ATO reduced spheroid viability and invasion while preserving the integrity and function of biomimetic endothelial vessels, suggesting limited toxicity toward healthy cells. Cell death was more pronounced in the spheroid core and intermediate regions, likely reflecting increased susceptibility to compounded metabolic and oxidative stress. In contrast, cells at the spheroid periphery remained viable but exhibited clear metabolic reprogramming, including a reduction in overall ATP production. Functionally, ATO significantly decreased the invaded area in both spheroid models and reduced migration distances, although a subset of cells retained the ability to migrate over longer distances within 72 hours of treatment.

At the invasive front of SKOV3 spheroids, ATO induced a shift in migration mode from individual to collective behavior, consistent with reduced energetic efficiency during early stages of treatment. At later time points, mitochondria in ATO-treated cells lost their polarized organization and instead appeared fragmented and perinuclear, accompanied by a more rounded cellular morphology. These observations suggest that while some cells initially retain migratory capacity, sustained OXPHOS inhibition disrupts mitochondrial organization and ultimately impairs continued migration.

We further demonstrate that OXPHOS inhibition modulates YAP1 signaling in a spatially restricted manner. ATO treatment reduced YAP1 expression in cells located at the spheroid periphery, indicating downregulation of the YAP/TAZ pathway in regions associated with active proliferation and invasion. This effect provides a potential mechanistic link between metabolic stress, reduced bioenergetic capacity, and impaired tumor progression. Consistent with this interpretation, mitochondrial network disruption in peripheral cells coincided with decreased migratory phenotypes, suggesting a coordinated response between metabolism, cytoskeletal dynamics, and mechanotransduction pathways.

Overall, our study highlights how metabolic heterogeneity shapes cellular behavior and therapeutic response and provides a more informative platform for evaluating metabolic therapies than conventional 2D systems. This approach is particularly relevant for assessing moderately potent OXPHOS inhibitors, such as metformin, which may exert subtle but spatially selective effects that are not detectable in bulk assays.

Several limitations should be noted. First, the cell lines used (SKOV3 and OVCAR5), although widely employed in ovarian cancer research, may not fully recapitulate the molecular features of HGSOC as defined by recent genomic studies. Future work incorporating additional patient-relevant models will be important to validate these findings. Second, the low aqueous solubility of ATO may have introduced local variability in drug distribution, as evidenced by crystal formation within the hydrogel. While ATO was delivered through diffusion rather than pre-incorporation into the matrix, it remains possible that similar heterogeneities in drug distribution could occur *in vivo*.

In summary, our findings demonstrate that OXPHOS inhibition with atovaquone selectively targets metabolically active and invasive regions of HGSOC tumors and limits invasion. We also provide a mechanistic hypothesis to these changes due to the suppression of YAP1 signaling in a spatially dependent manner. Future studies can further examine this potential mechanistic link and further leverage therapeutically. These results underscore the importance of spatially resolved models for understanding therapeutic response and highlight the potential of targeting metabolic plasticity as a strategy to constrain tumor progression while minimizing effects on healthy tissue.

## Materials and Methods

### Cell lines

OVCAR5 cells (ATCC) were maintained in RPMI 1640 (MT10040CV, Cellgro/Mediatech) supplemented with 20% FBS (MT35011CV), 1% Pen/strep (MT30002CI), 1% sodium pyruvate (MT25000CI), 1% sodium bicarbonate (MT25035CI), 0.5% of 45% glucose solution (in PBS), 1% HEPES 1M, and 0.1% insulin (I0516, Sigma). SKOV-3 (ATCC) were maintained in RPMI 1640 supplemented with 10% FBS, 1% sodium pyruvate, and 1% Pen/strep.

Human Umbilical Vein Endothelial cells (HUVEC, Lonza) were cultured in Lonza EGM-2-MV (CC-3202) in conventional cell culture flasks coated with fibronectin solution (Sigma F1141) diluted in PBS to achieve a concentration of 1 μg/cm^2^. Coating was performed for at least 1 h at 37°C and aspirated before seeding cells.

### Spheroid generation

Ovarian cancer cells were routinely trypsinized to generate spheroids using the hanging drop method. Briefly, cells were resuspended in corresponding media (see previous section) with 20% methylcellulose to form 25 µl droplets with 2,500 cells each. Droplets were incubated upside down for 48 h on the top of a 10 cm cell culture dish to allow the spheroids to form.

### Microfluidic device fabrication and setup

The device was fabricated using standard soft lithography methodology. Briefly, masks were designed using Illustrator software (Adobe) and used to generate two 500 and 750 μm-high SU-8 master molds (SU-8-100, MicroChem, Y13273) via conventional photolithography. Next, polydimethylsiloxane (PDMS, 10:1 base to curing agent ratio) was mixed, degassed, poured over each SU-8 wafer, and cured at 90°C for at least 2 h. The same PDMS mixture was injected into 25G needles and cured analogously to produce consistent 300 µm PDMS rods. The final microfluidic device consists of 2 layers of PDMS sandwiching one 300 µm PDMS rod and bonded to a MatTek dish glass plate (Diener Femto) to define the main chamber.

To enhance hydrogel adhesion to the device chamber, a three-step coating was performed through side-port injection and subsequent aspiration at room temperature: 1) 2% poly(ethyleneimine) (PEI, Sigma-Aldrich, 03880) in deionized (DI) water for 10 min, 2) 0.4% glutaraldehyde (GA, Sigma-Aldrich, G6257) in DI water for 15 min, 3) three DI water washes to clear any coating excess. To minimize evaporation, sacrificial phosphate-buffered saline (PBS, Fisher Scientific, BP3991, diluted to 1x with DI water) was added around the edges of the MatTek dish.

### Hydrogel mixing and cell seeding within hydrogels

Spheroids were harvested using a wide orifice tip and left to sediment in a 2 ml tube. A hydrogel mixture was mixed on ice, containing 4.5 mg/ml collagen (Corning), neutralized with NaOH 0.5M (Sigma-Aldrich, in DI), PBS 10X (Invitrogen) added to adjust osmolarity, and 30 μg/mL fibronectin (Sigma-Aldrich, F1141). Spheroids were pipetted into the mixture and carefully injected into the device chamber. After hydrogel polymerization, we removed the rod to reveal a tubular structure and pipetted the corresponding media.

HUVECs were routinely trypsinized and resuspended at 25,000 cells/µl, then injected into the tubular structure of an MPS containing a polymerized collagen hydrogel with the recipe mentioned above and incubated on either side for cells to attach and generate a fully coated structure to mimic the endothelial cell vessel.

### Treatment

Atovaquone (TCI America) was resuspended in DMSO at 60 mM and diluted in warm media before use at a final concentration of 30 µM. In the models, the atovaquone treatment was added through the lumen through multiple washing steps, to equilibrate the gel at the final atovaquone concentration and compensate for dilution within the chamber. DMSO was added as a vehicle control at 0.05%.

### Permeability studies

The barrier function of the endothelial cell vessel models treated with atovaquone was assessed by measuring the diffusion of a 1 µM solution of Texas Red dextran (70kDa, ThermoFisher Scientific, D1830) over 15 min^61^. 2 µl of this solution was added to each vessel via the small port. Dextran diffusion was tracked in a Leica Thunder Imager Live Cell inverted microscope (Buffalo Grove, Illinois). The permeability coefficients (P) were calculated from fluorescence intensity measurements (ImageJ) using equation 1^62^:

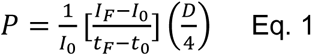

where *I_0_* is the total initial intensity outside the vessel, *I_F_* is the total intensity outside the vessel at 15 min, *t_0_* is the initial time point, *t_F_* is the final time point of 15 min, and *D* is the diameter of the vessel.

### Live cell staining and image analysis

Cells were stained with CellTracker (CMFDA, Thermo Fisher) at a dilution of 1:1000 per manufacturer’s instructions before forming spheroids to track growth over time. To assess viability, we stained devices with propidium iodide (PI, Thermo Fisher, 2.5 µg/ml in DMSO) and calcein AM (CAM, Thermo Fisher, 2 µg/ml in DMSO) for 30 min prior to imaging to ensure diffusion into the device. By complementing calcein AM with CMFDA, we found that overall spheroid staining was more effective. Conversely, PI diffusion into the spheroid and staining was not limited, therefore robustly producing a dead cell staining. Live-dead quantifications were done by subtracting PI staining from the total green staining (CAM + CMFDA). Quantification of the area of invasion was done by subtracting the total area occupied by the spheroid to the area of the spheroid core (center) at 72h. For the distance migrated, quantification was done by measuring the distance migrated by the cells from the outer edge of the spheroid and obtaining an average.

MitoTracker Green FM (Thermo Fisher) was used to stain mitochondria and determine their distribution around the nucleus. Briefly, a 1 mM stock solution in DMSO was diluted to 100 nM in relevant media with 1X SpyDye 650 (SpiroChrome, resuspended in anhydrous DMSO) to live-stain nuclei without cytotoxicity. The staining solution was added to spheroids after embedding and incubated at 37 °C for 1h, then media was refreshed prior to imaging. Cell aspect ratio was quantified using ImageJ by manually tracing cells in the periphery of the spheroid.

### Redox ratio imaging

Nicotinamide dinucleotide (NADH), nicotinamide dinucleotide phosphate (NADPH), and flavin adenine dinucleotide (FAD) were used to monitor the metabolic activity of the spheroids. The autofluorescence of these species enables a label-free per-pixel map of the oxidation–reduction state of a sample. Metabolic imaging was performed on a wide-field one-photon epifluorescence microscope (Leica Thunder 3D imager), and images were acquired sequentially for the same Field of View. NAD(P)H fluorescence was excited using an LED wavelength of 390 nm and an emission bandpass filter of 460/30 nm (30% LED power, 3 s). FAD fluorescence was excited using an LED wavelength of 475 nm and an emission bandpass filter of 535/30 nm (30% LED power, 5 s). The optical redox ratio was calculated as (FAD/ (FAD + NAD(P)H))^63^.

### Measurement of ATP levels

Cell Titer Glo 2.0 (G9242, Promega) was used to measure the levels of ATP in the spheroids. Briefly, spheroids (OVCAR5 and SKOV3) were embedded in a collagen hydrogel (recipe mentioned above) in a singular well of a TC-treated 96-well imaging plate (Corning, 353219) and treated with 30 µM of atovaquone and DMSO as a control in corresponding media. After 48 h of incubation, an equal volume of Cell Titer Glo 2.0 reagent was added to each well containing a single spheroid. After 1 h of incubation at room temperature, luminescence was read using a plate reader (Agilent, BioTek Synergy HTM Multimode Reader).

### Immunofluorescence

Washing buffer (PBS-0.1% Tween 80 (Sigma-Aldrich, P1754)) and blocking buffer (3% Bovine Serum Albumin (BSA, Sigma-Aldrich, A9056) in PBS-0.1%Tween 80) were made in advance and stored at 4°C until use. Unless specified otherwise, steps took place at room temperature. Cells were washed 3 times for 30 min between each step. Cells were fixed with 4% PFA for 15 min, then incubated with 0.2% Triton® X-100 (MP Biomedicals, 807426) for 30 min to permeabilize the sample. Next, devices were incubated with blocking buffer at 4°C overnight. To evaluate proliferation, cells were stained for Ki-67 (1:50, ThermoFisher, RM9106-S) or total YAP1 (1:100, Cell Signaling, 14074) overnight in blocking buffer and washed for 24 h afterward. Subsequently, we stained with a secondary antibody (1:1000, Thermo-Fisher A-11008) in blocking buffer overnight at 4°C. As cell indicators, we used Hoechst 33342 (Thermo Fisher, H1399, 1:1000 prepared per manufacturer’s instructions) and/or Phalloidin (1:400, Thermo Fisher T7471), both diluted in blocking buffer and incubated for at least 30 min. Devices were washed in PBS for a week before imaging. Z-stacks of the MPS were captured with a Leica Thunder 3D imager epifluorescence microscope. For ki67 staining, cells were manually counted to calculate % proliferation, whereas for YAP1 a plot profile of fluorescence intensity across a single confocal image slice was used as a measurement of expression.

### mRNA extraction and RT-qPCR

OVCAR5 spheroids were treated during 24h as described in the Materials and Methods section. Total RNA was extracted using TRIzol (#15596026, ThermoFisher) reagent according to the manufacturer’s instructions. For reverse transcription, 400 ng of RNA was used with the iScript Reverse Transcription Supermix (#1708840, Bio-Rad). Quantitative real-time PCR (qPCR) was performed using the Biosystems™ StepOnePlus™ Real-Time PCR System. Each PCR reaction contained 10 μL of SYBR™ Green PCR Master Mix (#4367659), 4.8 μL of nuclease-free water, 5 μL of cDNA template (diluted 10-fold from the reverse transcription reaction), and 0.2 μL of 20X primer stock solution (final concentration at 500 nM) of gene-specific primers (listed in Supplemental Table 1). Amplification was carried out under standard cycling conditions with an initial denaturation step at 95 °C for 3 min, followed by 40 cycles of denaturation at 95 °C for 10 s and annealing/extension at 60 °C for 30 s. Fluorescence data were collected at the end of each cycle. Relative gene expression levels were determined by normalizing the Ct values of target genes to the housekeeping gene RPL13A using the Pfaffl method.

### Western Blotting

Spheroids were embedded in collagen droplets in 6-well plates (approximately 12-15 spheroids per well in 100 μL droplets) and treated for 24h. At the end of the treatment, the culture medium was aspirated, and the collagen droplets were washed once with cold 1× PBS. The collagen droplets containing the spheroids were then transferred to 1.5 mL microcentrifuge tubes. To recover spheroids from the collagen matrix, collagen was enzymatically digested using collagenase (#17100017, Thermo Fisher). Briefly, collagenase was prepared in sterile PBS at a final concentration of 2 mg/mL. 1mL of collagenase solution was added to each tube, and samples were incubated at 37°C for 45 min under gentle agitation. During incubation, samples were gently pipetted every 10 min to facilitate collagen digestion and spheroid release while minimizing mechanical disruption. Once the collagen droplets were completely dissolved, the suspension containing the released spheroids was transferred to a fresh 1.5 mL microcentrifuge tube. 1mL of sterile PBS was added, and samples were centrifuged at 200 × g for 5 min. The supernatant was carefully removed. Spheroids were washed twice with sterile PBS to remove residual collagenase. For each wash, spheroids were gently resuspended in PBS using a wide-bore pipette tip and centrifuged again at 200 × g for 5 min.

After the final wash, spheroids were lysed in 50 μL of RIPA buffer (#89900, Thermo Fisher Scientific) supplemented with protease and phosphatase inhibitors. Protein concentrations were determined using a BCA assay, and equal amounts of protein (20 μg) were mixed with 4× Laemmli buffer (#1610747, Bio-Rad) and β-mercaptoethanol at a final concentration of 5%. Samples were denatured and separated on 4–20% precast polyacrylamide gels (#4561096, Bio-Rad) before transfer onto low-autofluorescence PVDF membranes (#1704275, Bio-Rad). Membranes were blocked for 1 h at room temperature using Intercept® (TBS) Protein-Free Blocking Buffer (#927-80001, LI-COR Biosciences) and subsequently incubated overnight at 4°C with primary antibodies against total YAP1 (#14074, Cell Signaling Technology), phospho-YAP1 (Ser94) (#254542, Abbiotec), and β-actin (sc-47778, Santa Cruz Biotechnology), diluted 1:1,000 for YAP1 antibodies and 1:10,000 for β-actin in Intercept® T20 Protein-Free Antibody Diluent (#927-85001, LI-COR Biosciences). Membranes were washed three times with TBST and incubated for 1 h at room temperature with IRDye® 680RD Goat Anti-Mouse IgG (#926-68070, LI-COR Biosciences) or IRDye® 800CW Goat Anti-Rabbit IgG secondary antibody (#926-32211, LI-COR Biosciences) diluted 1:10,000 in Intercept® T20 Antibody Diluent. Protein bands were visualized and quantified using the Odyssey CLx Imaging System (LI-COR Biosciences) and Image Studio software.

### Statistical Analyses

Data was analyzed using Graphpad Prism 11 software. All experiments were repeated at least three times as independent biological repeats for statistical significance. Results are presented as mean ± standard error. Statistical significance was set at p < 0.05. Comparisons between two samples were performed with an unpaired Student’s t-test assuming normal (Gaussian) distribution. Multiple comparisons by ANOVA were corrected using the Šidák test.

## Acknowledgements

The authors acknowledge funding support from the Office of Research on Women’s Health, Building Interdisciplinary Research Career in Women’s Health (BIRCWH) program, Centennial Scholars Foundation (UW-Madison), University of Wisconsin Carbone Cancer Center, and Diane Lindstrom Foundation.

## Author Contributions

Conceptualization, PMM, NB, MB, MVM; Methodology, PMM, NB, MB, EKL; Validation, PMM; Formal Analysis, PMM, NB, MB, MVM; Investigation, PMM; Resources, MVM, MP; Data curation, PMM, NB, MB, MVM; Writing-original draft, PMM, MVM, MB, YVN, ASPM; Visualization, PMM, MB, FRM, MVM; Supervision, MVM. All authors have read and approved the final draft.

