## Supplementary Figures for "Modeling the metabolic heterogeneity of high-grade serous ovarian cancer solid tumors in 3D Microphysiological systems"

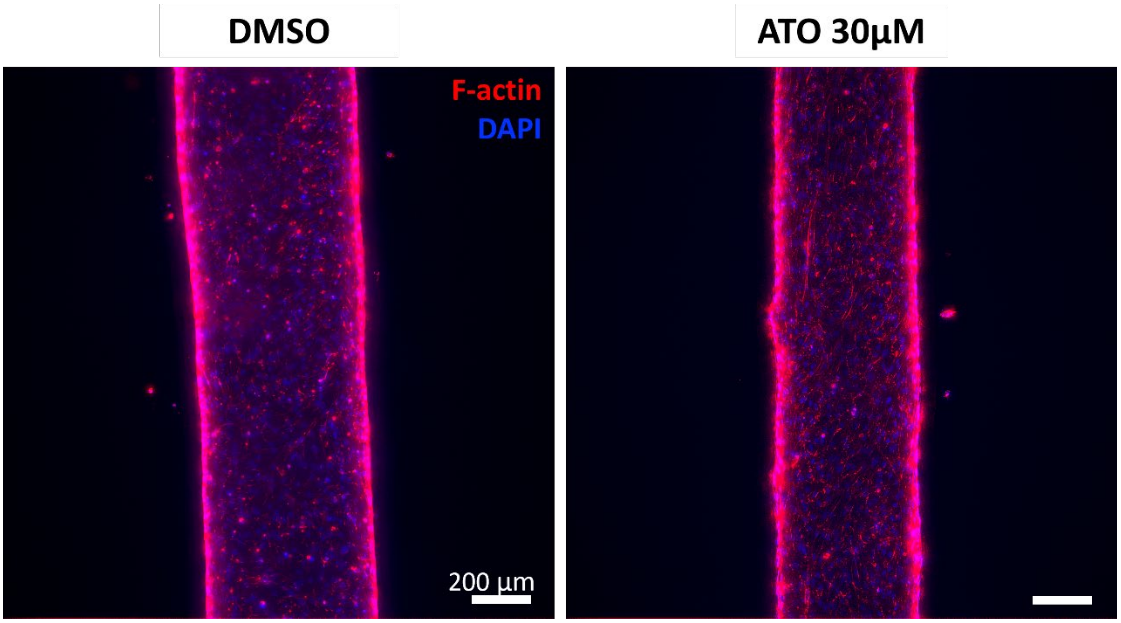


**Figure S1:** Fluorescent images of HUVEC vessels treated with ATO and DMSO stained with phalloidin to visualize the F-actin and DAPI.

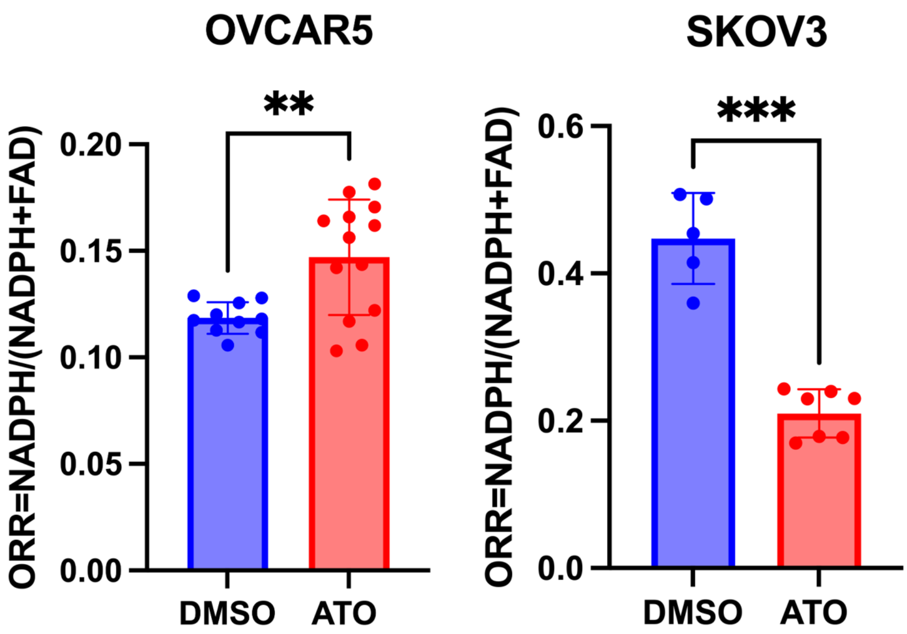


**Figure S2:** Optical redox ratio quantification for the OVCAR5 and SKOV3 spheroids for each treatment. Welch's t test was used. **p=0.003; p***<0.001; n=3 spheroids.

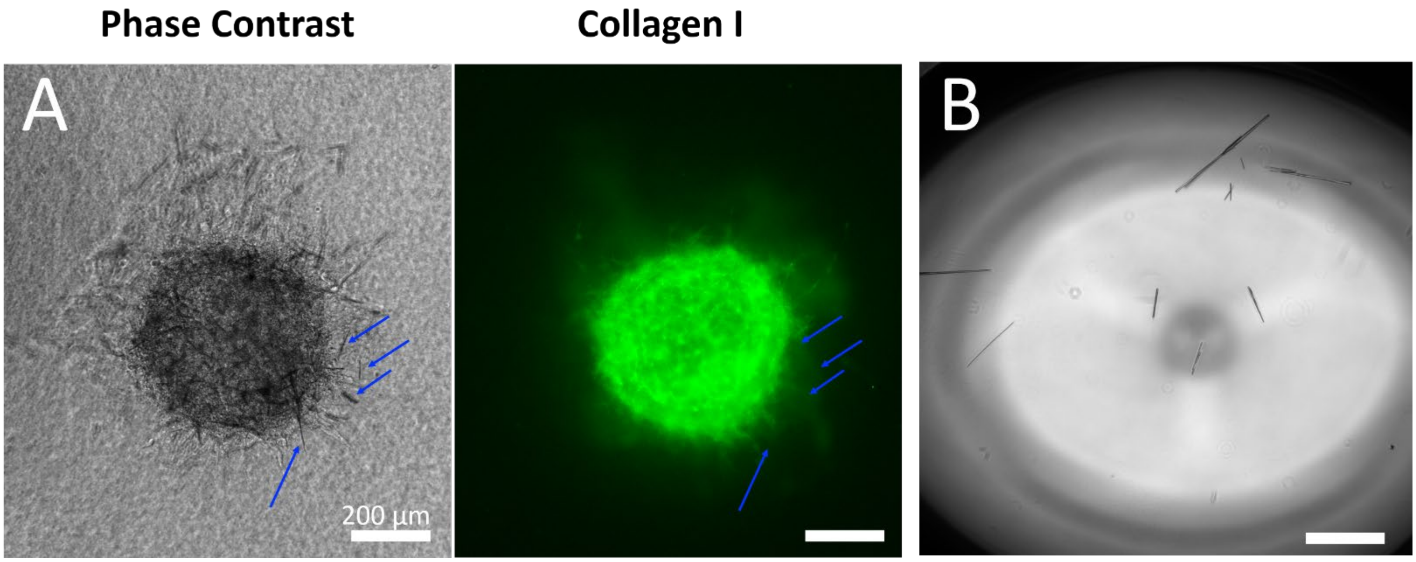


**Figure S3:** A) Phase contrast (left) and fluorescent image (right) of anti-collagen I immunostaining of SKOV3 spheroid treated with ATO to visualize ATO crystals surrounding the spheroid. Blue arrows indicate ATO crystals and are not observed in the anti-collagen I immunostaining. B) Phase contrast image of ATO crystals in media.

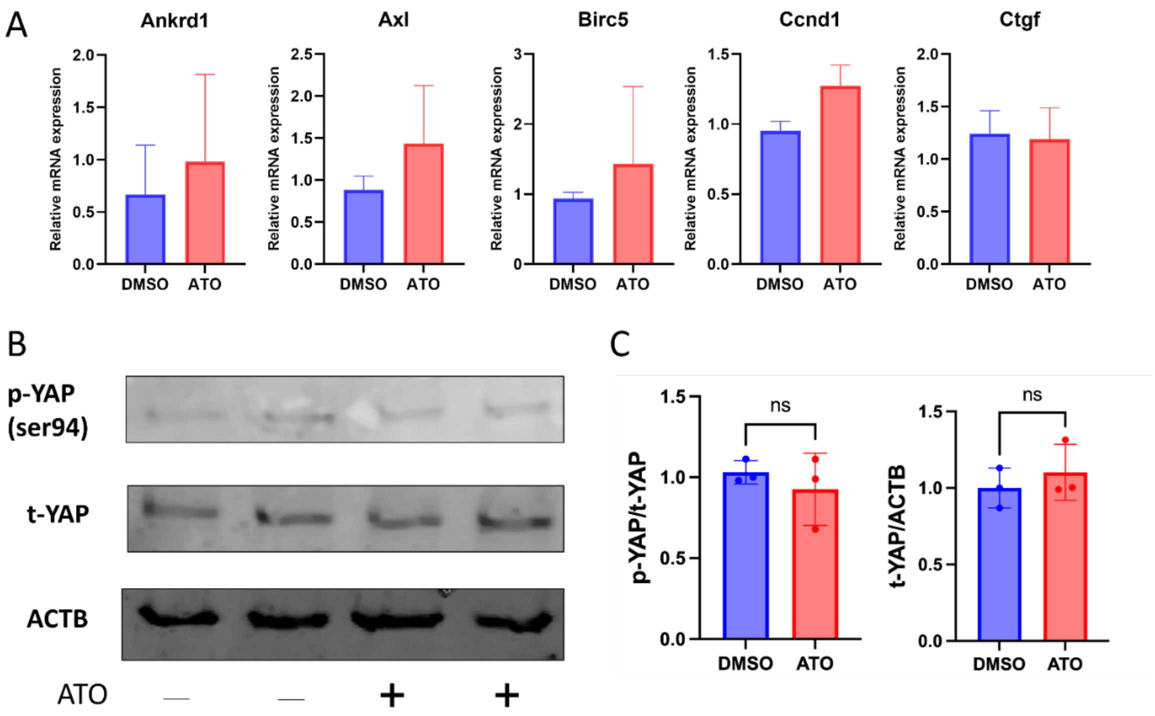


**Figure S4: Bulk analyses do not reveal significant modulation of YAP1 signaling following atovaquone treatment.** A) Expression of YAP1 target genes in OVCAR5 spheroids treated with atovaquone (ATO) or DMSO (vehicle control) for 24 h. Gene expression levels were normalized to the housekeeping gene RPL13A. B) Representative Western blot analysis of total YAP1 and phosphorylated AMPK (Ser94) following treatment with ATO or vehicle control. No statistically significant differences were observed between treatment groups ($p > 0.05$).

**Supplementary Table 1: qPCR Primer Sequences**

| Gene | Primer sequence |
|---|---|
| **Ankrd1 – F** | 5′-TCA ACG CCA AAG ACA GAG AAG-3′ |
| **Ankrd1 – R** | 5′-GTG TAG CAC CAG ATC CAT CG-3′ |
| **Axl – F** | 5′-CTG TGA AGA CGA TGA AGA TTG C-3′ |
| **Axl – R** | 5′-CAG AAC CCT GGA AAC AGA CA-3′ |
| **Birc5 – F** | 5′-CCA GTG TTT CTT CTG CTT CAA G-3′ |
| **Birc5 – R** | 5′-CAA ACT GCT TCT TGA CAG AAA GG-3′ |
| **Ccnd1 – F** | 5′-CCA GAG TGA TCA AGT GTG ACC-3′ |
| **Ccnd1 – R** | 5′-CGC AGG CTT GAC TCC AG-3′ |
| **Ctgf – F** | 5′-GCT CGG TAT GTC TTC ATG CTG-3′ |
| **Ctgf – R** | 5′-GAA GCT GAC CTG GAA GAG AAC-3′ |
| **Rpl13a – F** | 5′-CCT GGT GCT TGA TGG TCG AG-3′ |
| **Rpl13a – R** | 5′-CCT TCA CAG CGT ACG ACC AC-3′ |
